# Angelman syndrome patient neuron screen identifies a potent and selective clinical ASO targeting *UBE3A-ATS* with long lasting effect in cynomolgus monkey

**DOI:** 10.1101/2022.06.09.495066

**Authors:** Ravi Jagasia, Charlotte Bon, Soren V. Rasmussen, Solveig Badillo, Disa Tehler, Danièle Buchy, Marco Berrera, Megana Prasad, Marco Terrigno, Nikhil J. Pandya, Veronica Costa, Congwei Wang, Lykke Pedersen, Meghan T. Miller, Kamille Dumong Erichsen, Lars Joenson, Joerg Hipp, Azad Bonni, Lutz Müller, Annamaria Brändli-Baiocco, Thomas Kremer, Erich Koller, Marius C. Hoener

**Author notes:** These authors contributed equally to this manuscript (Ravi Jagasia, Charlotte Bon, Soren V. Rasmussen).

## Abstract

Angelman syndrome (AS) is a severe neurodevelopmental disorder caused by the loss of neuronal E3 ligase UBE3A with no available treatment. Restoring UBE3A levels via downregulation of the paternally cis-acting long non-coding antisense transcript (UBE3A-ATS) is a potential disease modifying. Developing molecules targeting human UBE3A-ATS is challenging because it is expressed only in neurons and lacks animal species sequence conservation. To overcome this, we performed a library screen of locked-nucleic acid (LNA)-modified antisense oligonucleotides (ASOs) on AS patient-derived neurons, identifying initial sequences. Further optimization led to the identification of the ASO, RO7248824, which selectively and potently reduces UBE3A-ATS, while concomitantly upregulating the UBE3A mRNA and protein. These properties held true in both human AS patient- and neurotypical-, as well as cynomolgus monkey-derived neurons. In vivo use of tool molecules in wild-type (WT) and AS Ube3am-/p+ mice, revealed a steep relationship between UBE3A-ATS knock-down and UBE3A mRNA/protein upregulation, whereby an almost 90% downregulation was needed to achieve a 50% upregulation, respectively. This relationship was confirmed in cynomolgus monkeys. Whereby, repeated lumbar intrathecal administrations of RO7248824 was well tolerated without adverse in-life effects or tissue pathology and produced a robust, long lasting (up to 3 months) paternal reactivation of UBE3A mRNA/protein across key monkey brain regions. Our results demonstrate that AS human pluripotent stem cell neurons serve as an excellent translational tool and furthermore LNA-modified ASOs exhibit excellent drug-like properties. Sustained efficacy translated to infrequent, intrathecal dosing and serves as the basis for the ongoing clinical development of RO7248824 for AS.

**Graphical Abstract:** Graphical abstract.
From AS patient blood to a neuronal screen, identifies clinical ASO with excellent *in vivo* properties.
(*1*) Patients were recruited. (*2*) Whereby blood was reprogrammed into hIPSC and subsequently differentiated into neurons. (*3*) ASOs were designed and screened on human neurons to downregulate the *UBE3A-ATS* likely via directed RNase H Cleavage of Nascent Transcripts. (*4a*) RO7248824 was identified that potently and selective reduces UBE3A-ATS, concomitantly with upregulating the *UBE3A* sense transcript and protein which was used for in vitro pk/pd. (*4b*) In parallel tool murine ASO were used demonstrate *in vivo* POC.(5) Pivotal nonhuman primate studies to monitor safety and predict the human dose. (6) *RO7248824 is in AS clinical trial*.

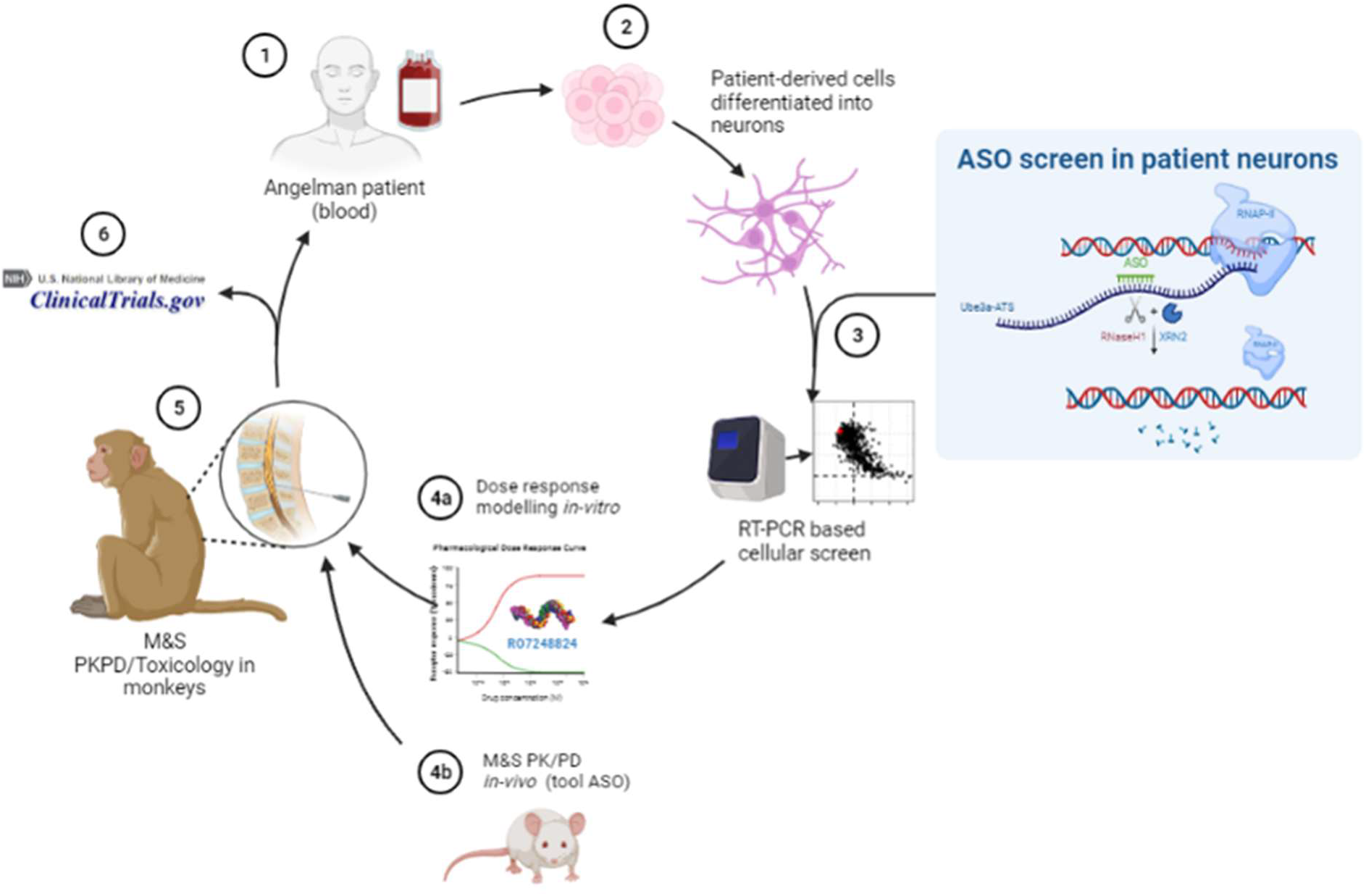

**One Sentence Summary:** From angelman syndrome human neuron screen to cynomolgus monkey proof of concept identifies the clinical molecule RO7248824

## Introduction

Angelman syndrome (AS) is a rare neurodevelopmental disorder that presents with severe symptoms including lack of speech, epilepsy, sleep disturbances and cognitive impairment (*1*). The genetic cause of the disorder is the loss of function of the maternal allele of the gene coding for Ubiquitin Protein Ligase E3A (UBE3A) (*2*–*4*). *UBE3A* is expressed biallelically in most tissues of the body except neurons. *UBE3A* regulation needs to be tightly controlled in CNS, since conversely to the loss in AS, overexpression contributes to the pathophysiology of another neurodevelopmental disorder, chromosome 15q11-q13 duplication syndrome (Dup15q; (*5*, *6*). Furthermore, two expressed copies of UBE3A may underlie increased susceptibility and severity of psychiatric illness in Prader Willi syndrome (*7*). The need for tight regulation of UBE3A may be due to the prescribed roles in fundamental neuronal and cellular function, including neuronal synaptic plasticity, proteasome and tRNA pathways and recently, in controlling viral derived GAG proteins (*8*–*11*); https://www.angelman-proteome-project.org/).

Controlling the regulation of UBE3A is achieved in large part via the expression of an antisense (ATS) transcript (*UBE3A-ATS*) only in neurons, which was discovered over two decades ago (*12*). Some years later, eloquent work demonstrated that *UBE3A-ATS* is an RNA polymerase transcript that represses Ube3a on the paternal chromosome, and this seminal work provided the rationale that deletion or knock-down of human *UBE3A-ATS* could provide a potential therapeutic strategy for AS, including the use of antisense oligonucleotides (ASOs) (*13*). Subsequently, it was demonstrated that genetically depleting *Ube3a-ATS* in mice could activate expression of Ube3a from the paternal chromosome, and could revert disease-related symptoms in the AS mouse (*14*). Since these discoveries of the function of *Ube3a-ATS*, multiple therapeutic approaches using ASOs have demonstrated ‘’proof of concept’’ by interfering with this process, and in turn reinstating the expression from the paternal *UBE3A* allele to levels comparable to maternal *UBE3A* (*9*, *14*–*17*). Of note, ASOs have also been shown to enable modulation of active transcription by targeting the nascent RNA including murine *Ube3a-ATS* (*15*, *18*). In addition, other modalities such as small molecules and genomic medicines targeting genetic insertion of a transcriptional termination cassette have been reported (*9*, *14*–*17*). The promise of these therapeutic approaches have been supported by the finding that reinstating Ube3a expression after postnatal development in AS mouse models, either using a tamoxifen-inducible cre-system or via ASOs, demonstrates partial rescue of phenotypes (*17*, *19*, *20*).

These studies with AS animal models have been instrumental to support therapeutic strategies that focus on increasing neuronal UBE3A expression levels by targeting *UBE3A-ATS* transcription (*19*). However, lack of sequence conservation of *UBE3A-ATS* between rodents vs. humans and limited understanding of the neuronal regulation of the paternal allele across species in the postnatal brain has hampered translational research. To overcome these challenges, here we present an *in vitro* screening strategy leveraging human AS patient neurons that recapitulate the *UBE3A-ATS*-dependent paternal allele silencing to identify potent *UBE3A-ATS*-targeting LNA ASOs. RO7248824 was identified using this strategy, has excellent *in vitro* and *in vivo* potency, selectivity and is well tolerated in mice and monkeys. RO7248824 produces a long-lasting reduction in *UBE3A-ATS* in monkey brains and a concomitant upregulation of paternal *UBE3A* mRNA and proteins. Data was used to derive a translational pharmacokinetic/ pharmacodynamic model and enabled human interventional studies.

Our data highlight the benefits of screening on human patient neurons, and incorporating LNA chemistry into oligonucleotide therapeutics to maximize effect duration for the treatment of AS.

## Results

### Tiling screen identifies ASO targeting UBE3A-ATS to unsilence the paternal UBE3A allele with human/cynomolgus monkey cross reactivity

To model a physiologically relevant neuronal network which comprises both excitatory and inhibitory neuronal connectivity, we leveraged a protocol that differentiates Neuronal Precursor Cells (NPCs) derived from human Induced Pluripotent Stem Cells (hiPSC) into a mixture of both glutamatergic and GABAergic neurons (*9*, *21*, *22*). The patient neurons develop significant synaptic maturation as evidenced by synchronous network activity by day 42 of neuronal differentiation (*9*, *21*, *22*). We had previously described and characterized hiPSC lines for AS individuals harboring deletions of the 15q11-13 locus and one from an individual harboring a point mutation in UBE3A (*8*, *9*). Over the course of neuronal differentiation from hiPSC to NPC to neurons there was an increase in the expression of the *UBE3A-ATS* in both control and AS patient lines with a concomitant decrease of the *UBE3A* mRNA transcript (Figure 1B). The difference of *UBE3A* mRNA expression in AS neurons plateaued by day 42 in culture and was confirmed using immunohistochemistry where *UBE3A* levels were very low in the soma of TAU positive neurons (Figure 1B, 1C). We concluded that our AS hiPSC-derived neuronal model recapitulates the loss of *UBE3A* in disease. We reasoned that this model would be well-suited for screening of RNaseH recruiting LNA/DNA ASO gapmers to target and down-regulate human *UBE3A-ATS*. All ASOs tested had phosphorothioate backbones and were composed of DNAs in the gap and LNAs in the flanks. The kinetics of *UBE3A* imprinting in AS neurons, plateauing at day 42 of neuronal differentiation, guided the treatment paradigm that resulted in the development of a screen to identify specific, safe, and efficacious ASOs targeting human *UBE3A-ATS* (Figure 1E).

**Figure 1:**
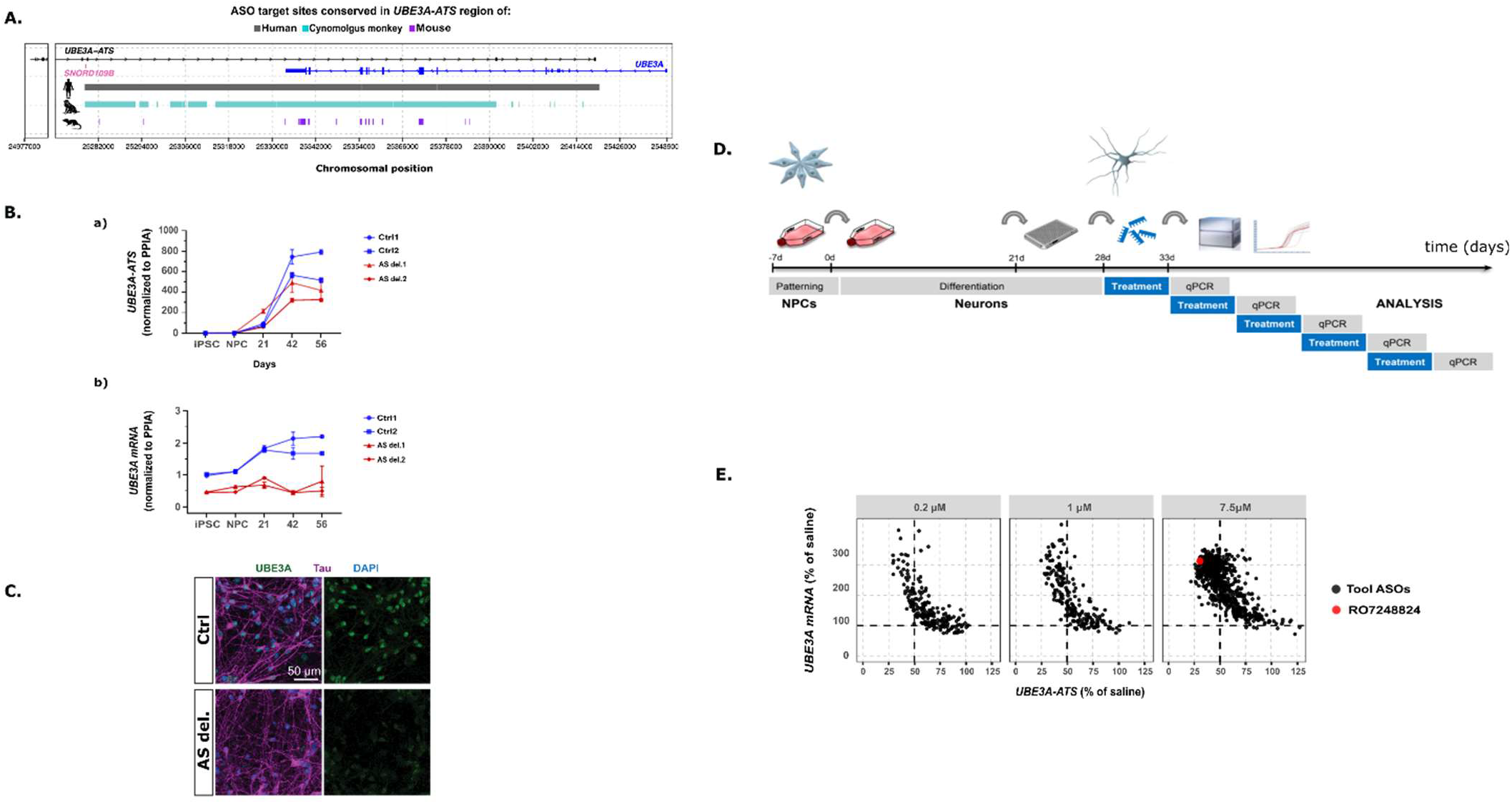
Human Angelman syndrome patient derived neuron screen identifies UBE3A-ATS targeting sequence RO7248824 that upregulates the UBE3A sense transcript. (A) Human genetic locus of UBE3A mRNA and UBE3A-ATS in the overlapp–25,439,008). Colored bars denote the start position of 20-mer ASOs designed to target this region in human (gray) if their target site is present in the corresponding region of cynomologous monkey (cyan) or mouse (purple). (B) qPCR analysis for UBE3A-ATS (top) or UBE3A sense transcript (bottom) normalized to housekeeping gene PPIA in control and AS iPSC, neuronal precursor cells (NPCs), and at Day 21, Day 42 and Day 56 of differentiation. Control lines 1302 and 902 and AS del. 1501 and 1301 lines were used. Each data point represents independent neuronal differentiations. n = 4.PCR (C) Immunostaining of UBE3A (green) antibody along with Tau (magenta), in control 1302 and AS del. 1501 lines at Day 56 of differentiation (Scale bar: 50 μm).IHC (D) Scheme for screening AS hIPSC derived neurons with ASOs targeting UBE3A-ATS. (E) Results of the ASO screen correlating UBE3A mRNA (y-axis) and UBE3A-ATS (x-axis) in hIPSC-derived AS neurons after incubation with ASOs. Levels at indicated concentrations measured by RT-qPCR relative to GAPDH and normalized to PBS treated cells from two separate ASO screens in biological duplicates with CV < 30%.The effect of RO72488 is highlighted in red at 7.5 μM (triangle).

Human neurons were initially differentiated from NPCs derived from iPSCs, and then subsequently replated in a 96 well plate. The entire process took approximately five weeks (Figure 1D). We designed approximately 2’500 RNase H recruiting ASOs targeting *UBE3A-ATS* across a region spanning from the most distal snoRNA in 15q11-q13 (*SNORD109B*) to the 3’ end (chr15:25,280,119-25,414,623) (Figure 1A, 1E). Since the species conservation is lacking between rodents and humans, we looked for sequences in *UBE3A-ATS* that were conserved between human and cynomolgus monkey, that passed safety and specificity criteria based on predictive algorithms, to allow for later characterization of ASO safety, efficacy, and translational PK/PD modeling in non-human primates (NHP) (Figure 1A). ASOs were screened for their potential to upregulate *UBE3A* mRNA using AS patient iPSC-derived neurons with the assumption that increasing *UBE3A* would represent unsilencing of the paternal allele (Figure 1E). This effort led to the identification of sequence “hotspots” on *UBE3A-ATS* in regions both upstream of and overlapping with the *UBE3A* gene body that, when targeted by ASOs, led to strong increase in *UBE3A* mRNA passing our cutoff for efficacy of 2-fold upregulation (Figure 1E). These data demonstrate a good correlation between *UBE3A-ATS* knockdown and *UBE3A* upregulation revealing that 50% *UBE3A-ATS* downregulation unsilenced the paternal locus in the vast majority of cases, whereas for some ASOs 30% knock-down was enough to achieve unsilencing (Figure 1E). Based on efficacy, off target predictions and safety algrorithms we selected the molecule RO7248824 depicted in red for further characterization (Figure 1E).

### RO7248824 potently and selectively unsilenced the paternal UBE3A allele in human and cynomolgus neurons

RO7248824 targets a sequence homologous to both human and cynomolgus monkeys (Figure 2A). Immunocytochemical analysis demonstrated that MAP2 positive soma of AS patient iPSC-derived neurons show reduced UBE3A immunoreactivity which is restored by RO7248824 (Figure 2B). Dose-response treatment of AS neurons with RO7248824 revealed nanomolar potency for *UBE3A-ATS* (IC_50_ = 26.3 nM), UBE3A mRNA upregulation (EC_50_ = 15.4 nM) and UBE3A protein upregulation (EC_50_ = 24.8 nM) (Figure 2C, 2D). RO7248824 showed similar potencies in neurons derived from neurotypical iPSCs as well as in neurons derived from cynomolgus iPSCs, consistent with biallelic *UBE3A* expression (Figure 2C, 2D). These cross primate *in vitro* neuronal results suggest that RO7248824 upregulation of *UBE3A* mRNA translates to increased UBE3A protein levels in neurons derived from both AS patient iPSCs, neurotypical iPSCs and cynomolgus iPSCs (Figure 2B, 2D).

**Figure 2:**
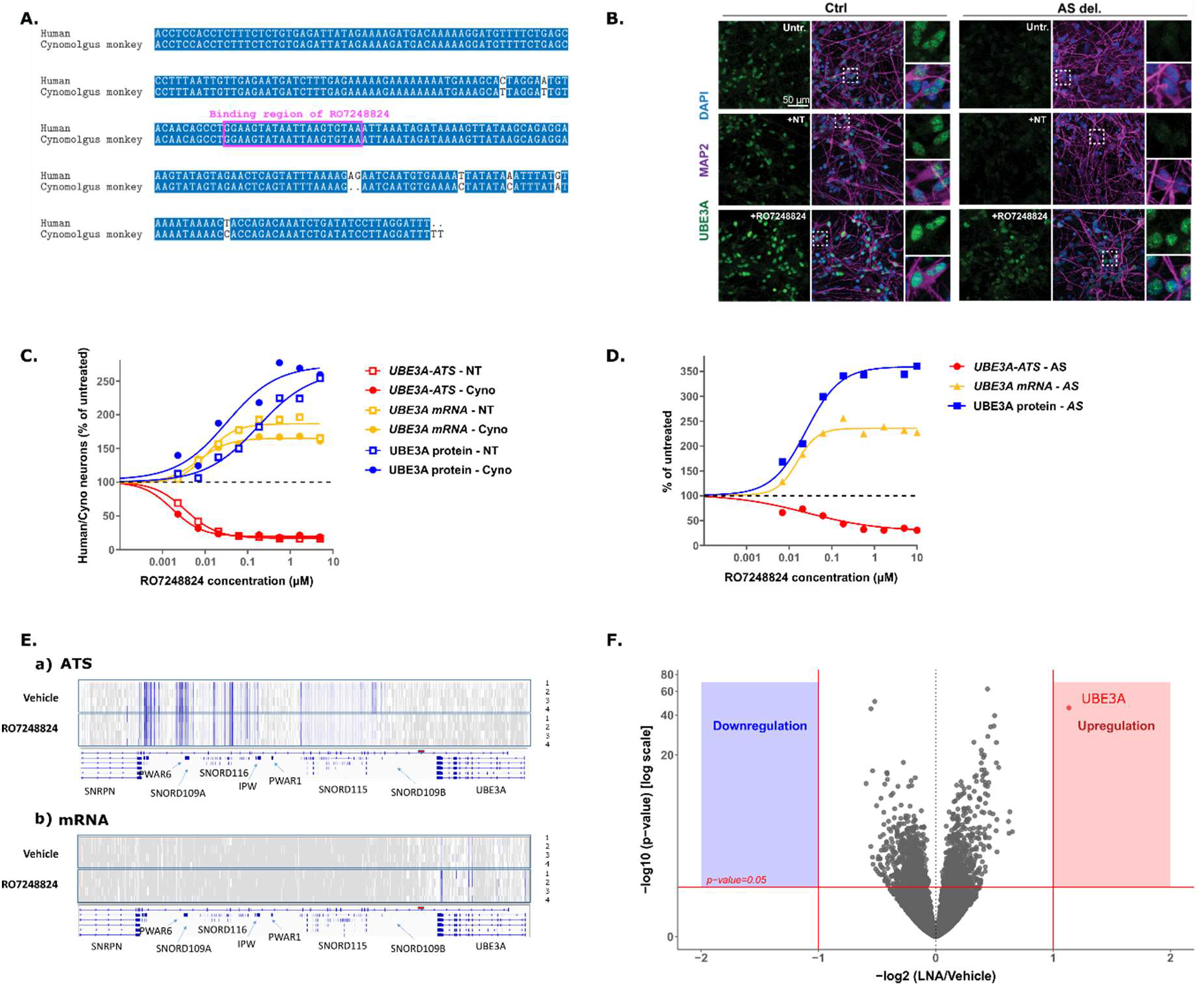
RO7248824 is a highly potent and selective molecule targeting and reactivating the UBE3A locus on human and cynomolgus monkey neurons. (A)Sequence alignment of the region surrounding the binding site of RO7248824 (magenta) in human and cynomologous monkey corresponding to the genomic positions chr15:25,308,946-25,309,225 (hg38) and chr7:3,314,713-3,314,992 (macFas5). (B) Immunostaining of UBE3A (green) antibody along with MAP2 (magenta), in control 1302 and AS del. 1501 lines. After 56 days of differentiation, cell lines were left untreated (Untr.) or treated with a non-targeting ASO or RO7248824 1 μM for seven days (Scale bar: 50 μm). (C) Dose-response curves to assess pharmacological properties of RO7248824 *in vitro* using human (derived from non-diseased subjects, circles) and cynomolgus monkey (wild-type animals, boxes) iPSC-derived neurons (*UBE3A-ATS*, in red; *UBE3A* mRNA, in yellow; UBE3A protein, in blue). (D) Dose-response curves to assess pharmacological properties of RO7248824 *in vitro* using iPSC-derived neurons from derived from an Angelman Syndrome subject (*UBE3A-ATS* RNA, in red; *UBE3A* mRNA, in yellow; UBE3A protein, in blue). (E) Single reads from the stranded RNAseq on the *UBE3A* gene locus and the up-stream loci in control and RO7248824-treated iPSC-derived neurons. Coverage is represented in blue for the positive chromosomal strand that encodes *UBE3A-ATS* (a) and negative strand that encodes *UBE3A* mRNA (b). (F) Selective UBE3A upregulation in RO7248824-treated iPSC-derived neurons. Volcano plot showing the genes dysregulated by RO7248824 treatment. The red lines show the thresholds applied for log2FoldChange and adjusted p-value to identify differentially expressed genes. A single transcript, UBE3A (red dot), was significantly upregulated upon treatment.

To identify any potential off-target RNA downregulated by RO7248824, we performed whole transcriptome sequencing in AS-derived neurons after 48h incubation with 30 uM of RO7248824. Differential gene expression analysis comparing RO7248824 and vehicle treated cells demonstrated a single transcript, *UBE3A*, to be upregulated and no other transcripts significantly (adj. p < 0.05) downregulated by 2-fold or more (Figure 2F) which is remarkable since the tested concentration was 1000-fold higher than the *in vitro* efficacious dose. Since *UBE3A-ATS* lies at the 3’-most end of the *SNHG14* transcript, which also encodes multiple other small RNAs, we next examined the whole transcriptome data for downregulation of other RNAs within *SNHG14* by RO7248824. Consistent with the proposed mechanism of action of RO7248824 to target the nascent RNA of actively transcribed *UBE3A-ATS*, the expression of *SNHG14* most proximal to the ASO target site (see Figure 1A), such as portions of the RNA that include *SNORD109B* and *SNORD115*, was reduced upon treatment (Figure 2E, Top panel). However, the majority of the other portions of *SNHG14*, such as the majority of the *SNORD115* cluster, the *SNORD116* cluster, *PWAR, SNRPN*, etc. showed similar levels of expression as seen in the control samples (Figure 2E, Top panel).

### Tool ASO targeting murine *Ube3a-ATS* to characterize PK/PD relationship in AS model mice

Before embarking on studies using NHPs, we wanted to demonstrate *in vivo* “proof of concept’’ and in this context understand the relationships between the *UBE3A-ATS* knock-down and the UBE3A protein expression to support pivotal translational studies. To that end, we screened LNA/DNA gapmers targeting murine *Ube3a-ATS* in mouse primary cortical cultures to identify a tool molecule for *in vivo* studies as previously described (*20*). The screen identified both RTR26183 and RTR26235 as potent activators of *Ube3a* mRNA in primary cortical neuron cultures prepared from Ube3a^m-/p+^ pups (EC_50_ = 23 nM) as well as WT littermates (EC_50_ = 38 nM) (*23*). The targeting sequence of RTR26183 with respect to mouse genomic position is depicted in Figure 3H.

**Figure 3:**
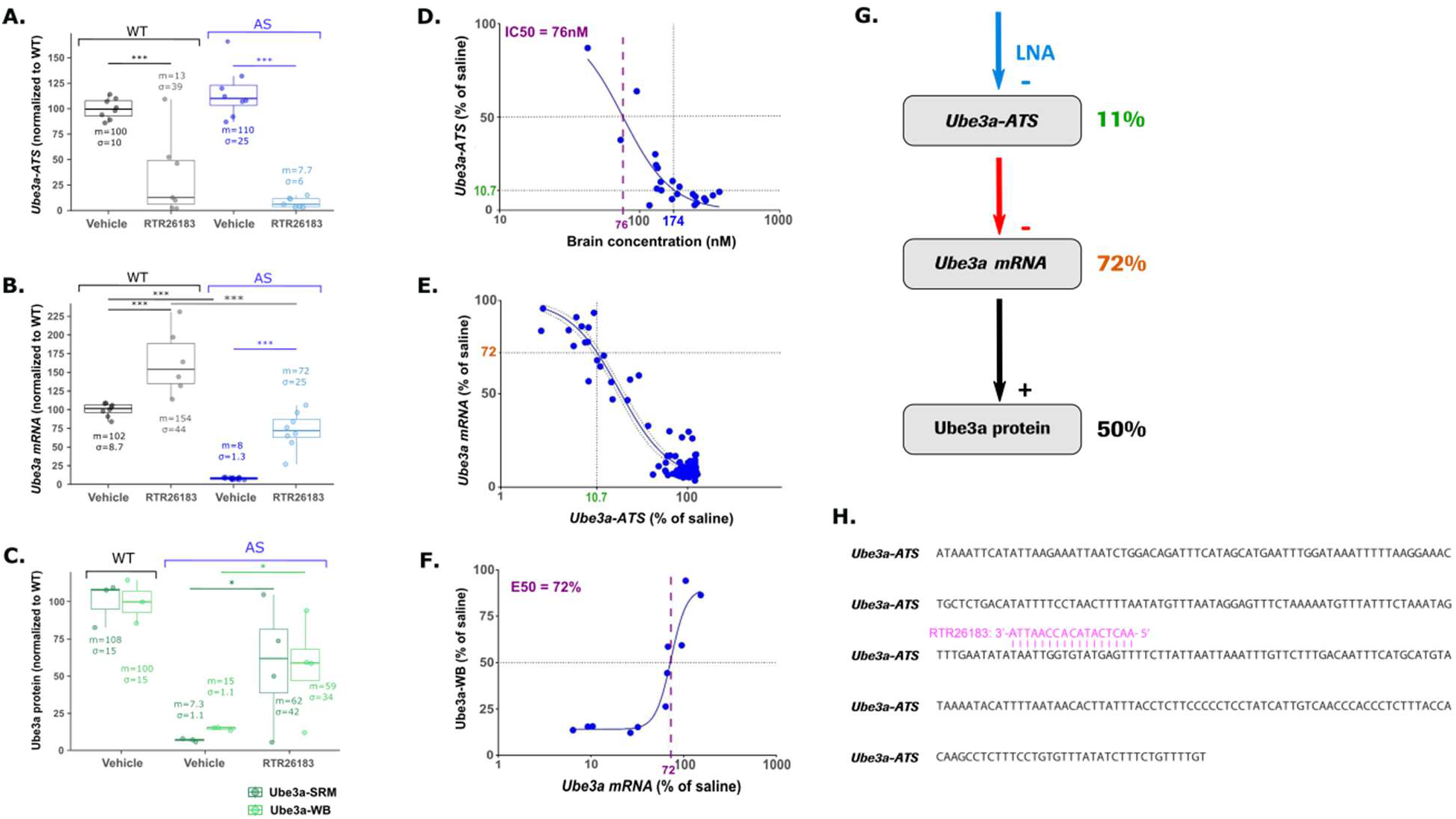
Single dose of a potent mouse tool molecule, RTR26183, exhibits sustained reactivation of paternal locus in both WT and AS model. (*A)Ube3a-ATS* knock-down with oligo RTR26183 in both WT and AS. In vehicle-treated animals, in both AS and WT, the *Ube3a-ATS* (a) is expressed and suppressed upon oligo treatment. (B)The *Ube3a* mRNA increases with oligo RTR26183, to a level of 75% and 150% of control in WT and AS mice respectively. (C) Ube3a protein is increase with oligo RTR26183 to a level of in AS mice. (D) RTR26183 concentration dependent knock-down of the *Ube3a-ATS*. The blue line represents the Emax model where at 76 nM the *Ube3a-ATS* is down to 50% of the level in saline treated animals. (E) PKPD relationship between the knock-down of the *Ube3a-ATS* and the *Ube3a* mRNA elevation. The blue line represents the Emax model and the dotted lines represent the 95% confidence interval. *Ube3a* mRNA is elevated to 50% when the *Ube3a-ATS* is knocked-down about 18.5%. (F) PKPD relationship between the *Ube3a* mRNA elevation and the Ube3a protein. The blue line represents the Emax model, when the *Ube3a* mRNA is elevated at 72% the protein is translated at 50%. Data from experiments conducted with RTR26183 and RTR26235 have been analyzed together since the relationship between these two pharmacological biomarkers are independent of the compound. (G) Quantitative relationship between *Ube3a-ATS* knock-down, *Ube3a* mRNA and protein expression in mice. To elevate the protein to a 50% level versus saline treated group, it requires a very high knock-down of the *Ube3a-ATS*, almost 90%. All biomarkers are represented at steady-state where the tool LNA knock-downs the *Ube3a-ATS* (blue arrow) which suppresses the silencing mechanism (red arrow). As the *Ube3a* mRNA is transcribed the protein is then translated (black arrow). (H) Mouse sequence of *Ube3a-ATS* surrounding the binding site of RTR26183 (magenta) corresponding to the genomic position chr7:58,989,211-58,989,527 (mm39). Median (m) and standard deviation (σ) values are indicated for each group, and stars above horizontal lines indicate significant levels of corresponding groups comparison: *** for adjusted p-value<0.001 and * for adjusted p-value<0.05 (t-tests).

A single 150 ug ICV dose of RTR26183 and RTR26235 was well tolerated and sufficient to knock-down *Ube3a-ATS* by more than 90% in both WT and AS Ube3a^m-/p+^ mice in adult mice in brain regions (Figure 3A). *Ube3a* mRNA was upregulated by 55% in WT mice (Figure 3B), while *Ube3a* mRNA was paternally unsilenced to 76% of WT levels in Ube3a^m-/p+^ mice (Figure 3B). To confirm treatment effect translated to protein changes, Ube3a protein levels were quantified in AS mouse tissue via western blotting and mass-spectrometry. Both methods confirm a significant increase of Ube3a protein expression in the treatment group, in line with previous results (Figure 3C) (*17*, *20*).

A PK/PD model was derived to link ASO brain exposure to *Ube3a-ATS*, *Ube3a* mRNA and protein levels from different brain regions including hippocampus, striatum and cortex at 14 days post injection in Ube3a^m-/p+^ mice (Figure 3D-F). RTR26183 had *in vivo* potency on *Ube3a-ATS* knock-down in the nanomolar range (~75 nM, Figure 3D). To investigate the relationship between *Ube3a* mRNA and its protein, the data from *in vivo* experiments with RTR26235 was added. Indeed, the relationship between the mRNA and the protein is believed to be compound independent. Interestingly, correlating the degree of *Ube3a-ATS* knock-down with paternal *Ube3a* mRNA and protein elevation, we found that almost 90% *Ube3a-ATS* knock-down was required to increase *Ube3a* mRNA by 72% which translated to 50% upregulation of protein (Figure 3D-F). This steep relationship was not completely in line with iPSC neuron *in vitro* data and reveals that a large knock-down of the *Ube3a-ATS* silencing mechanism is needed to allow protein reexpression in WT mice. This data on one side brought confidence in the *in vivo* mode-of-action and inspired the study design for RO7248824 in pivotal NHP studies needed for human dose prediction and subsequent translation.

### RO7248824 distributes throughout the brain causing a widespread and sustained unsilencing of the paternal *UBE3A* locus in NHPs

Since the *in vitro* profile of RO7248824 on patient, neurotypical and cyno neurons was both potent and selective and *in vivo* potency was demonstrated with the tool molecule in mice, the molecule was evaluated in cynomolgus monkeys using IT delivery as the route of administration to assess pharmacological properties, safety, and to develop a human PK/PD model. In brief, 3 male cynomolgus monkeys per group and time point were dosed by an intrathecal administration (ported catheter) (Figure 4A). Either saline or a single dose of 24 mg RO7248824 or two doses of 16 mg separated by two weeks and subgroups were sacrificed at different time points spanning from day 8 to day 85 (animals dosed twice on days 1 and 15 were sacrificed on day 29) (Figure 4A). At the selected time points of sacrifice, brains were harvested for both measuring RO7248824 concentrations and the three PD markers *UBE3A-ATS* RNA, *UBE3A* mRNA and UBE3A protein in order to estimate brain uptake and clearance of RO7248824 and understand the dynamics of target engagement as well as relationship to paternal unsilencing (Figure 4A). Levels were analyzed in multiple brain regions thought to be neurologically relevant for AS clinical phenotypes including cortex (frontal, occipital, parietal, temporal), hippocampus and cerebellum along with spinal cord regions (lumbar, thoracic, cervical).

**Figure 4.**
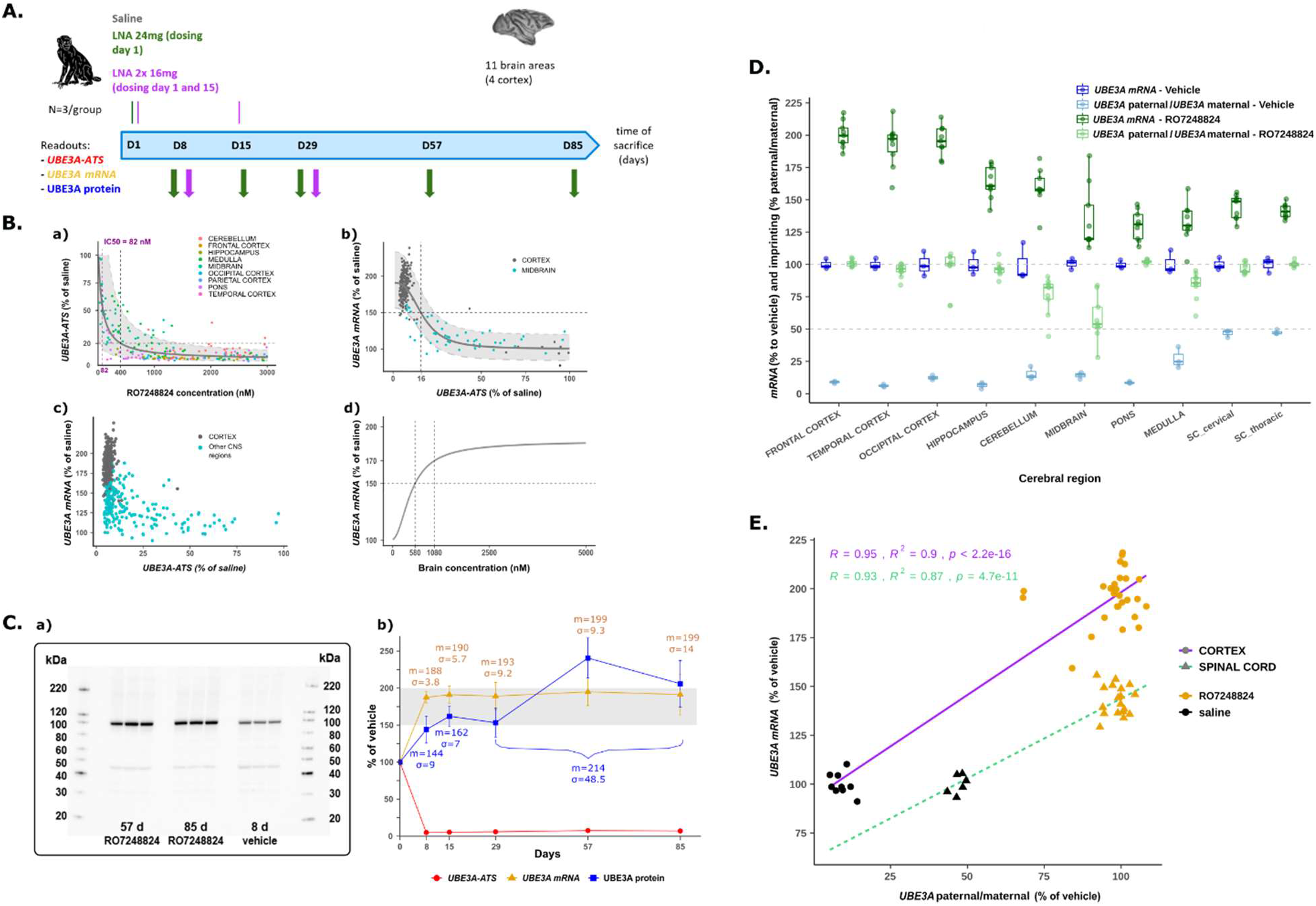
Single dose of RO7248824 has a long duration of action on paternal UBE3A reactivation in NHP brains after IT delivery. (A) Schematic on NHP study. Either saline, a single dose of 24 mg or 2 doses of 16 mg 2 weeks apart was doses intrathecally. Sacrifice was 8, 15, 29, 57, and 85 days after the last dose. 9 brain regions were analyzed for UBE3A-ATS and UBE3A mRNA and UBE3A protein concomitantly with ASO concentration. (B) a) Exposure dependent reduction of UBE3A-ATS. Measured values are shown as dots that are colored coded depending on the tissues. The grey plain line represents the median model fit with the variability depicted as a shadow area (90% confidence interval around the median). At 395 nM the UBE3A-ATS is reduced by about 80%. b) UBE3A mRNA expression is doubled upon full UBE3A-ATS knock-down. UBE3A-ATS knock-down is followed by UBE3A mRNA elevation if the knock-down is sufficiently high. When the UBE3A-ATS is reduced to 16% UBE3A mRNA is elevated to about 150%. Measured values are shown as dots that are colored coded depending on the tissues. The grey line represents the median model fit with the variability depicted as a shadow area (90% confidence interval around the median). c) The expression of the UBE3A mRNA is doubled in the cortex regions but to a much lesser extent in other brain regions despite the UBE3A-ATS being knocked-down (90%). d) Steep relationship between brain LNA concentration and UBE3A mRNA expression. 1080 nM is required in brain tissue to ensure 170% mRNA upregulation. Plain line represents the median prediction of UBE3A mRNA upregulation versus concentration. (C) a) Western Blots, b) Long lasting knock-down and high level of UBE3A upregulation in cortical regions of cynomolgus monkey (WB, 24 mg SD). Squares indicate median values (also indicated with m), and standard deviations (σ) are shown with vertical segments. For protein values, we show median values and standard deviations for three time points grouped together: days 29, 57, 85 because the effect is delayed for protein. (D) Paternal gene expression relative to maternal gene expression in brain tissue following treatment with saline or RO7248824. (E) Correlation between UBE3A protein (SRM) and ratio between paternal/maternal UBE3A mRNA expression in cortex (magenta) and midbrain (green) after saline or RO7248824 treatment.

Intrathecal administration of 24 mg RO7248824 showed high brain uptake (> 5’000 nM seven days post injection, which is approximately 50-fold higher than the *in vitro* potency on upregulation) in most brain regions with the highest exposure observed in cortical regions and the lowest in midbrain (Figure 4B). The tissue half-life was determined to range from 6-7 weeks in brain regions (data not shown). The combination of high brain uptake and long half-life are potentially ideal properties to ensure infrequent intrathecal dosing in clinics. RO7248824 also distributed in plasma and from there to peripheral organs such as the kidney and liver (data not shown). Liver exposure was in the same range as seen for hippocampus regions but the kinetics was faster with a decreased half-life in the liver of approximately 1.5 to 2 weeks. Exposure in the kidney was about five times higher than in liver, highlighting the main elimination pathway with intrathecal dosing. In this context, it was encouraging to recognize the absence of any clinical or histopathological evidence for toxicity to the liver or kidney (data not shown). Transient lack of coordination was observed in most animals administered RO7248824 at 24 mg. These changes correlated with transient neurological observations noted at 2 and/or 6 hrs post dose which included decreased proprioceptive hindlimb positioning, decreased or absent tactile and/or visual hindlimb placing reaction, decreased or absent hindlimb flexor reflex and/or decreased muscle tone (data not shown). These types of reactions are rather typical for intrathecally dosed ASOs and are generally not adverse (*24*). In terms of histopathological analysis, test item related findings of neuronal vacuolation in the nervous system, vacuolated macrophages in spinal cord, injection site and lymph nodes, and basophilic granules in the kidneys were not considered adverse based on their low severity and lack of associated degenerative/inflammatory changes or clinical pathological changes (assessing renal function). All of these safety-directed observations were considered consistent with those observed after administration of single-stranded oligonucleotides (*25*, *26*).

In all brain regions analyzed, RO7248824 exposure correlated with reduction of *UBE3A-ATS* (Figure 4B). Remarkably, a knock-down of > 95% was observed in cortical areas relative to the saline controls which was maintained up to day 85 (last sacrifice). In deeper brain structures, such as the midbrain, the exposure and knock-down was more variable while not as pronounced, resulting in 75% reduction at day 7 (Figure 4B). The calculated *in vivo* potency was similar in all tissues, and we found that *UBE3A-ATS* is reduced by 80% at 400 nM, which was approximately 20 fold less potent than the potency from *in vitro* human neurons (Figure 4B).

In cortical areas, knock-down of *UBE3A-ATS* translated to upregulation of almost 200% of *UBE3A* mRNA expression versus saline, and sustained throughout the 85 days of the experiment. 200% is the theoretical maximum, since in the NHP brains the contribution from the maternal allele should be present from baseline (Figure 4B panels a-c). In the midbrain, the PD effects (*UBE3A-ATS* and *UBE3A* mRNA) were less pronounced compared to cortex regions, likely due to lower exposure and faster clearance in this region (Figure 4B panel b). In other deeper brain regions such as medulla and particular pons the upregulation of *UBE3A* mRNA was less pronounced and more variable despite high exposure and strong decreases in *UBE3A-ATS*, especially shortly after intrathecal injection on day 8 and 15 (Figure 4B panel c). A direct response model, correlating the *UBE3A-ATS* knock-down to *UBE3A mRNA* upregulation in cortex and midbrain, similar to the mouse, revealed that a knock-down of at least 84% was needed to achieve a 50% increase, respectively (Figure 4B panel d). Using this PD direct response model together with the *UBE3A-ATS* versus concentration model, we derived the relationship between concentration and *UBE3A* mRNA expression to be an EC_50_ of 580 nM (Figure 4B).

From day 8 to day 85 the protein increase correlated well with the *UBE3A* mRNA expression in cortical regions (data not shown). *UBE3A-ATS* levels were reduced to about 6% of controls at all time points. *UBE3A* mRNA expression in cortex was increased to roughly 190% throughout the duration of the study (Figure 4C panel b). Both western blot and SRM analysis, revealed that after some delay, the protein level was doubled in cortical regions and this level was sustained during the whole experiment (Figure 4C panel a-b). We concluded that due to the favorable PK properties (a high uptake and long half-life), a single administration of RO7248824 is able to fully reinstate paternally-derived UBE3A protein in NHP cortical regions for at least 12 weeks.

To summarize, both, in-life and detailed histopathological analyses of the response to treatment with RO7248824 triggered initiation of further formal short-term and long-term GLP toxicity studies and clinical development. These findings were confirmed in formal short-term GLP toxicity studies conducted thereafter in support of dosing in AS patients.

### RO7248824 equalizes paternal and maternal UBE3A expression across most brain regions

The relative increase in *UBE3A*mRNA in deeper brain regions and spinal cord tissues was more variable than in cortical regions (Figure 4B). Specifically, besides high exposure and strong knockdown of *UBE3A-ATS*, the relative increase in UBE3A mRNA increase was far less pronounced in the spinal cord than cortical regions. One explanation could be the differential degree of paternal allele silencing across discrete brain regions. To assess the relationship between paternal silencing and UBE3A protein upregulation, we made use of a subgroup of animals that are heterozygous *UBE3A* SNP carriers (n = 12 in our sample from both saline and RO treated groups at day 15 and 29 post injection, Chr7: 3.339.686C → A) identified by sequencing. We developed allele-specific *UBE3A* mRNA digital droplet PCR assay to assess contribution of the paternal versus maternal allele to overall *UBE3A* mRNA expression in respective brain tissues (Figure 4D).

Allele-specific *UBE3A* mRNA expression analysis in various brain regions showed differences in baseline imprinting levels, ranging from paternal/maternal ratio of approximately 0.05 in cortical regions to 0.5 in spinal cord tissues (Figure 4D and E). Notably, tissue with the highest degree of imprinting (lowest paternal/maternal ratio in the saline group) showed higher abundance of UBE3A mRNA (Figure 4D). In all cortical brain regions of the treated animals, an equal expression of paternally vs. maternally derived *UBE3A* was observed (ratio ~1), which indicates complete unsilencing of the paternal allele. In the cerebellum, midbrain and pons, the absolute increase in *UBE3A* mRNA was less pronounced than in cortical regions (+/− 14%, N=17 samples from 9 animals, Figure 4D) and allele specific quantification showed an almost complete unsilencing (1.02 +/− 0.017, N =17 samples from 9 animals) (Figure 2B). There was a significant correlation between the efficiency of paternal unsilencing and UBE3A mRNA. In deeper brain tissues, such as hindbrain tissues (medulla, cerebellum) and spinal cord tissues (thoracic and cervical) the allele specific unsilencing (Figure 4E), demonstrated equal *UBE3A* mRNA expression of paternal and maternal alleles.

We assessed the relationship between degree of imprinting at baseline and increase in *UBE3A* mRNA in regions with differential degree of imprinting such as cortex, medulla and spinal cord. All regions showed strong knock-down of *UBE3A-ATS* but varying degree of mRNA increase. Correlation analysis suggests that the degree of imprinting determines the extent of a UBE3A upregulation (Figure 4E). In the cortex, an equal expression of paternally vs. maternally derived *UBE3A* (ratio ~1) led to almost doubling of *UBE3A* mRNA whilst in the spinal cord the lower ratio resulted in far less effect size.

### PKPD modeling of the *UBE3A* LNA ASO to support human dose selection

A PKPD model was derived from the pharmacokinetics of the ASO concentrations and the time course from target engagement to paternal locus unsilencing (relationship between *UBE3A-ATS*, *UBE3A* mRNA and UBE3A protein) in NHP brain tissue. RO7248824 showed brain uptake well above the *in vivo* IC50 on *UBE3A-ATS* and EC_50_ on *UBE3A* mRNA (> 5 μM seven days post injection) in most brain regions with the highest exposure in cortical regions and lowest in midbrain. The half-life of RO7248824 was estimated around 6-7 weeks but varied across brain regions (data not shown). Assuming that the relationship between *UBE3A-ATS* and brain concentration is similar across brain regions, we found that a direct response model fitted best the data (Figure 4B). Potency was measured in the nanomolar range and the slope of the response is steep (hill coefficient >1). The *UBE3A* mRNA response is also described by a direct response model, linking remaining *UBE3A-ATS* (%) to the *UBE3A* mRNA elevation (%). The estimation of the IC50 is about 16%, meaning that the knock-down needs to be higher than 84% to produce half maximal response in the *UBE3A* mRNA elevation (Figure 4B). The finding is in line with the WT and AS mouse model, where a knock-down of 90% is needed to reinstate *UBE3A* mRNA to 70% (Figure 3). The link between brain ASO concentration and *UBE3A* mRNA elevation was derived from the relationships between *UBE3A* mRNA/-ATS and *UBE3A-ATS/* ASO concentration (Figure 4B panel C). The protein expression followed the *UBE3A* mRNA expression but with some delay (Figure 4E). Therefore, the protein response is described with a turnover model where the *UBE3A* mRNA and protein levels have a 1:1 relationship under steady-state conditions. The half-life of the protein was estimated at about 13 days which is in line with protein half-life in the brain (*27*). Despite a steep relationship between ASO concentration/ *UBE3A-ATS* knock-down and resulting *UBE3A* paternal transcription, the combination of a high uptake and long ASO half-life explains why the UBE3A protein stays elevated for months after a single intrathecal dose. With a 24 mg dose level, the steady state at the protein level is achieved with an interval of dosing every 16 weeks in monkeys. This very long interval between dosing can be attributed to the combination of both the long ASO- and protein half-lives. This model provided the justification for the human dose selection in AS patients.

## Discussion

The fact that all AS patients carry at least one intact copy of paternal *UBE3A*, which is silenced by the long non-coding RNA, *UBE3A-ATS*, in almost all neurons, brings considerable therapeutic opportunity. Since there is substantial molecular evolutionary divergence between humans and mice within the *UBE3A-ATS* sequence, we developed an AS patient derived neuronal cell model to screen for gapmer ASOs with LNA chemistry in the flanks, targeting *UBE3A-ATS*. This led to the identification of potent and selective human/cynomolgus monkey cross reactive ASOs, including RO724884. Before conducting pivotal NHP studies, we tested tool ASOs in a mouse model of AS. Our AS patient-derived neurons served as an excellent disease relevant cellular model for screening and enabled detailed characterization of the relationship between the three key events culminating in paternal allele unsilencing: defining the relationship between ASO concentration, *UBE3A-ATS* knock-down and concomitant upregulation of *UBE3A* mRNA and protein. This mechanism of action was observed *in vitro* in iPSC-derived in both neurotypical and derived human AS neuron, cynomolgus monkey neurons, as well as *in vivo*, after ICV injection of a tool molecule in WT and an AS mouse model, and after intrathecal administration in NHPs. Remarkably, RO724884 in juvenile to pubertal NHP resulted in sustained and almost complete neuronal unsilencing of the paternal *UBE3A* locus in brain regions likely important to AS clinical presentation including the forebrain for cognitive and speech impairment and cerebellum for motor disability (*28*, *29*). Even 3 months after the injection there is still almost doubling of the *UBE3A* mRNA and protein in various cortical regions. The data from the NHP study were used to derive a translational PK/PD model, suggesting an acceptable dosing frequency in patients. This work combined with the desirable safety profile culminated in RO7248824 entering clinical trials for AS (clinicaltrials.gov identifier: NCT04428281).

Mouse reactive *Ube3a-ATS* ASOs had been previously identified which led to both knock-down of the *Ube3a-ATS* and a partial restoration of Ube3a protein (*14*, *17, 20*). These ASOs resulted in a reversal of deficits in an AS mouse model linked to the disease including synaptic plasticity and cognition (*17*, *20*). In addition, human specific ASOs revert functional and molecular changes in patient neurons (*9*, *15*). In a recent study, human/cynomolgus monkey cross reactive ASOs targeting the *UBE3A-ATS* revealed moderate knock-down of the *UBE3A-ATS* but failed to upregulate *UBE3A* mRNA and protein levels in non-AS neurons and NHP brains (*30*). It was hypothesized to be due to potential feedback regulatory mechanisms that hinder upregulation of *UBE3A* mRNA and protein in a non-disease context (*30*). Here, by generating data across three different species, specifically human derived neurons, NHPs and mice, we provide substantial evidence that potent knock-down of *UBE3A-ATS* enables unsilencing of paternal *UBE3A* expression with maximal UBE3A protein upregulation even in a “healthy context”, with no evidence for maternal allele compensation. The discrepancy between these results might be explained in part by molecule properties (eg. binding affinities) or alternatively the region it is targeting on the *UBE3A-ATS* locus. For example, RO7248824 targets an intronic sequence which may be more effective in terminating transcription downstream of ASO-mediated cleavage of nascent transcripts; which is considered crucial for the proposed silencing mechanism of *UBE3A-ATS* (*31*). Furthermore, the precise location relative to other genomic elements (i.e. splice sites, protein binding sites and others) may influence the ability of an ASO to activate *UBE3A*, since these other elements may also influence transcriptional termination (*15*, *32*). Another not well understood mechanism, albeit very important for modeling and human dose predictions, is the steep relationship between *UBE3A-ATS* lowering and unsilencing in both mouse and cynomolgus monkey brains but not in the human neurons. Again, here, future work is needed to elucidate the underlying mechanisms, for example the ASO concentration we measured was from bulk brain tissue and not necessarily from the active compartment in the cell, specifically the nucleus. Single cell level nuclear resolution analysis, comparing ASO-mediated *UBE3A-ATS* cleavage, transcriptional termination, and *UBE3A* transcription might give some answers and may reveal how the ASO may impact on the so-called “RNA polymerase collision hypothesis” at this locus (*13*–*15*, *33*).

Intrathecal injections are relatively invasive compared with orally delivered medications, thus identification of molecules with long duration of action are greatly needed to develop drugs with infrequent administration. Compared to earlier generations of ASO chemistries, LNA modified ASOs are well-known for their improved affinity to target RNA, this may well be a property potentially resulting in the longer duration of action (*34*). RO7248824 showed a desirable pharmacokinetic profile, whereby the ASO produced a notably extended duration of effect of paternal unsilencing in monkeys, up to 3 months after intrathecal dosing, in key disease brain regions. Due to the invasive nature of intrathecal administration, requiring an in-patient procedure, infrequent dosing is of critical importance. Similar to other oligonucleotides tested in the NHP and humans, RO7248824 did not distribute to all tissues equally in NHP brains (*35*). We found ASO concentration and efficiency of paternal unsilencing was highest in cortical regions with less in the cerebellar nuclei, ventral midbrain, pons, and medulla. Few data have been published on oligonucleotide brain pharmacokinetic in patients after intrathecal administration but it seems that concentrations in the micromolar range are achievable (*35*, *36*). It remains to be seen if the intrathecal administration of RO724884 will be as efficient in patients as it was in the NHPs.

Importantly, concentration alone cannot explain the variability of protein expression across brain regions since the spinal cord had high ASO distribution whilst less efficiently leading to paternal allele unsilencing. Future work aimed at more homogenous delivery of ASO payloads may provide further insights by for example crossing over the blood brain barrier by hijacking transport systems such as the transferrin receptor or by the recently reported DNA/RNA cholesteralized molecules (*37*, *38*).

Several challenges for clinical development of any *UBE3A-ATS* ASO therapeutic remain, including understanding the brain distribution of intrathecally injected ASOs in human, and importantly biomarkers to measure pharmacodynamic responses on a molecular and circuit level. Whether UBE3A can be measured in CSF, and more specifically if treatment response could be quantified, remains to be determined. Alternatively, proteins downstream of UBE3A in neurons measurable in body fluids may allow monitoring proximal target engagement after restoring UBE3A for example as recently reported for PEG10, TKT and likely others (*8*, *9*); and see: https://www.angelman-proteome-project.org/). In addition, EEG (excess delta-band power) is emerging as an UBE3A-disease relevant biomarker of abnormal brain function in AS and rodent models that has been linked to symptom severity in natural history data (*39*–*41*). It will be important to understand the relationship between these candidate biomarkers to clinical severity and outcome measures both in the natural history and in response to treatments.

Collectively, our data support the clinical development of RO7248824 in patients with Angelman syndrome (clinicaltrials.gov; NCT04428281).

## Materials and Methods

### Human neuronal cell culture, Cynomolgus monkey iPSC derived NPC culture and LNA treatment

NPCs derived from neurotypical hIPSC lines 1302 a-C (ND 140321, L500) and CRA902 and AS patient iPSC lines (1501a-A(A) (EdS150402) and CRA1301 were previously described and characterized (Pandya et al, 2022; Pandya et al, 2021). The neuronal differentiation protocol was previously described (*21*, *22*). In brief, hIPSCs derived NPCs were dissociated with Trypsin-EDTA 0.05% (Thermo Scientific), plated on polyornithine-/laminin-coated (PL) flasks/dishes at 10k cells/cm2 in SFA medium, and cultured for one week with medium replacement after 3-4 days. The resultant progenitors were dissociated with trypsin/EDTA 0.05%, plated on PL flasks/dishes at 50k cells/cm2 in BGAA medium and differentiated for 6 weeks with medium replacement every 3-4 days. *UBE3A-ATS-* or non-targeting-LNA (NT) was incubated at indicated concentration from day 0, unless otherwise noted, and LNA was maintained in the cell culture media.

NPCs derived from cynomolgus macaque (Cips_BC671A_NSC line generated at Roche) were differentiated in NPC differentiation medium (*42*). Differentiating neurons were incubated at indicated concentration from day 15 with ASO. Cells were lysed for qRT-PCR analysis at day 20.

### Generation of human/cynomolgus cross specific ASO and screen

Based on previous characterization of the regulatory relationship between *UBE3A-ATS* and *UBE3A* mRNA (Meng et al., 2014) we designed about 2500 RNase H recruiting ASOs targeting different sites of the *UBE3A-ATS* transcript between the its last annotated snoRNA SNORD109B and the 3’end of *UBE3A-ATS* (chr15:25,280,119+ to chr15:25,414,623+) as previously described by (*43*). To allow for later *in vivo* characterization of ASOs in NHP all ASOs were targeting sequences conserved between human and cynomolgus monkey (Figure 1A). Conservation was evaluated by aligning the target sites of all possible 20-mer oligonucleotides designed against the human region spanning the last annotated snoRNA and the 3’end of *UBE3A* mRNA and the region spanning the end of *UBE3A* mRNA to the 3’end of *UBE3A-ATS* to the corresponding regions in mouse and cynomologous monkey (Figure 1A). The region surrounding the target site of RO7248824 was specifically extracted from human and cynomologous monkey sequences and aligned with the msa package in R using the ClustalW method Figure 2A.

In the course of optimizing for ASO screening (Figure 1D-F) as an effort to make the neuronal cultures more homogenous and reproducible, AS patient 21 day differentiated cultures were replated. On day 21 medium was aspirated and cells were washed with PBS-/-,prior to addition of 6 ml pre-warmed Accutase/ T75 flask, and incubated for 5 min at room temperature. Cells were resuspended in 6 ml of NDM+BGAA+Laminin medium and centrifuged in a 15 ml Falcon tube at 1’100 rpm for 4 min. Supernatant was removed and cells were resuspend in 10 ml of NDM+BGAA+laminin medium, counted and seeded in the same medium additioned with ROCK inhibitor at the concentration of 400’000 cells/cm2 in 96 well plates. At day 28 ASOs were added (final concentration of 0.2, 1 or 7.5 μM) and RNA was harvested (RNeasy mini kit [Qiagen, Cat#: 74106]) after 5 days of treatment. The RNA levels of *UBE3A-ATS* (Assay ID: Hs01372957_m1, FAM, Thermo Fisher). *UBE3A* mRNA (Assay ID: Hs00166580_m1, FAM, Thermo Fisher) and GAPDH (housekeeping reference gene; Assay ID: 1_Hu GAPDH VIC 4325792, Thermo Fisher) were evaluated in duplicates by qRT-PCR according to the manufacturer’s instructions. The expression of *UBE3A* mRNA and *UBE3A-ATS* were normalized to GAPDH expression from the same well and shown relative to the mean of the PBS samples represented at each qPCR plate. Values with a coefficient of variance below 30% for UBE3A were plotted in Figure 1E and F.

### Generation of mouse specific tool compound

ASOs (N=192) were designed and synthesized as previously described (*43*). All ASOs were complementary to the minus strand of chromosome 7 in a 34 Kb region (chr7: 59307929 – 59341825) downstream of UBE3A which is partly overlapping with the annotated transcript, Snhg14 (*Ube3a-ATS*). The ASOs were screened at a single concentration for their ability to knockdown *Ube3-ATS* in embryonic (E14-15) mouse primary cortical cultures (data not shown). The potency of the most efficacious compounds was determined by generating concentration response curves in primary cortical cultures prepared from Ube3a^m-/p+^ P1 pubs measuring both *Ube3a-ATS* and Ube3a mRNA expression by qPCR. This identified the ASO, RTR26183, with fully modified phosphorothioate (PS) backbone and the nucleotide sequence, AACTcatacaccaatTA capital letters are locked nucleic acid (LNA) modifications and lower case letters are DNA. All LNA-C nucleotides contain the 5-methylcytosine modification.

#### Neuronal cellular readouts: mRNA and protein

##### 1) qPCR analysis for determination of UBE3A mRNA and UBE3A-ATS in vitro

RNA purification from both IPSC and primary mouse cortical cultures were done using RNeasy 96 Kit (Qiagen) according to the manufacturer protocol. The purified RNA was diluted in water to a concentration suitable for the one-step RT-PCR reaction (XLT-Onestep, Quanta). The RT-PCR reaction was carried out on a ViiA 7 384-well Real-Time PCR System (Thermo-Fisher) running a standard 40 cycle program according to the manufacturer protocol. For quantification of *UBE3A* mRNA, *UBE3A-ATS*, GAPDH in human iPSCs we used Taqman assays Hs00166580_m1 (UBE3A), Hs01372957_m1 (*UBE3A-ATS*) and Hs 4325792 (GAPDH) all from ThermoFisher. For quantification of *UBE3A* mRNA, *UBE3A-ATS* and Gapdh in mouse primary cortical cultures we used Mm02580987_m1 (UBE3A), Mm03455899_m1 (*UBE3A-ATS*) and Mm99999915_g1 (GAPDH) all from ThermoFisher.

All reactions were carried out in duplicates and on each QPCR plate a standard curve was included with 2-fold serial dilution of RNA from PBS treated control wells. Using the QuantStudioTM software (Applied Biosystems) RNA quantities were calculated based on the standard curve. Quantities were then normalized to the calculated quantity for the housekeeping gene (GAPDH) in the same well. Hence, Relative Target Quantity = QUANTITY_target gene/ QUANTITY_housekeeping gene. The RNA knock-down level was calculated for each well by dividing with the median Quantity of all PBS-treated wells (N=14) on the same plate. Normalized Target Quantity = (Relative Target Quantity / [median] Relative Target quantity_PBS_Wells). This way expression level data of *UBE3A* mRNA and *UBE3A-ATS* are shown as percentage of PBS-treated wells.

For concentration response experiments, cultures and oligonucleotide addition were carried out as described above. Compounds were added to the cells in an 8 or 10-step, ½-log serial dilution. Concentration response curves were fitted using Graphpad Prism 7, using a four parameter sigmoidal dose-response model. Expression values from PBS treated cells were included in the curve fitting, by setting the PBS treatment to fixed concentration of a full log10 below the lowest tested ASO concentration.

##### 2) AlphaLisa assay for determination of UBE3A protein levels in vitro

hiPSC derived neurons were cultured in 96 well plates. Medium was aspirated, wash carefully twice with PBS, add 50 μl of lysis buffer/well, incubate on ice for 5 min, mix the samples by pipetting to ensure good lysis of the cells, proceed with analysis or store at −80 °C.

Lysis buffer: in 20 mM Tris HCl pH 8, 137 mM NaCl, 10% Glycerol Acros Ref.184695000, 1% NP-40 Sigma Ref.74385, 2 mM EDTA Gibco Ref.15575-038, phosphatase inhibitors (PhosSTOP, Roche Ref.04906837001), protease inhibitors cocktail (Complete, Roche Ref 46931132001)

The standard curve was used for normalization of sample values and was included in each assay plate. The curve included 11 concentrations with 1:2 dilution steps ranging from 1.25 to 0.0012 μg/ml. For the standard curve the recombinant protein E6AP/UBE3A, Boston Biochem Cat.#E3-230 was used.

For each sample the assay reagent mix was prepared, containing: 5 μl of sample (cell lysate or standard curve), 10 μl of biotin conjugated antibody UBE3A-1/10a (in house AB) to a final concentration of 1 nM, 10 μl of AlphaLisa beads-conjugated antibody UBE3A-1/4b (in house ab) to a final concentration of 10 μl/ml, 25 μl of Streptavidin AlphaLisa donor beads to a final concentration of 40 μg/ml. Samples were prepared in a 384 well plate (OptiPlate, Perkin Elmer Cat. #6007290). After preparation, the plates were centrifuged at 1’200 rpm for 2 min at RT, the plates sealed and incubated for 30 min at RT. The plates were read using the instrument SpectraMax (Molecular Devices). Specific details of the stocks of conjugated antibodies used were reported in each original data file.

##### 3) Whole transcriptome sequencing from hiPSC neurons

Total RNA was isolated from iPSC-derived neurons in two batches using the RNEasy Mini Kit (Qiagen) according to the manufacturer’s protocol. An on-column DNAse digestion was included to deplete genomic DNA.350 ng total RNA per sample was used as input for library preparation. Ribosomal RNA was depleted using the Ribo-Zero Gold rRNA Removal Kit (Illumina). rRNA-depleted RNA was further processed into sequencing libraries using the TruSeq Stranded Total RNA kit (Illumina) according to manufacturer’s instructions. Cluster generation was performed on the cBot instrument and paired-end sequencing reads were subsequently generated on a HiSeq4000 instrument.

To estimate gene expression levels, paired-end RNASeq reads were mapped onto the human genome (hg19) by using the short read aligner GSNAP version 2017-05-08 (Wu et al., 2005; Wu et al., 2010) with default alignment parameters and allowing the program to detect novel splice. Mapped reads for all RefSeq transcript variants of a gene were combined into a single value, read counts per gene, by applying SAMtools version 1.5 (Li et al., 2009) and customized inhouse tools. Subsequently, read counts were normalized by sequencing library size and gene length according to Mortazavi et al. (Mortazavi et al., 2008), denoted as RPKMs (number of mapped reads per kilobase transcript per million sequenced reads). Prior to differential gene expression analysis (LNA vs. vehicle, n = 4 each), a negative binomial regression model was derived to correct for potential confounding factors with the inclusion of the following covariates: lane, percent mapped reads, RNA Integrity Number and an estimated IPS differentiation to neuron score based on the derived Fantom5 primary cells genesets information. The implementation was conducted in R (version 3.4.0) using the DESeq2 package (*44*) (version 1.16.0).

#### Mouse Intra-cerebroventricular (ICV) injections

In both WT and AS Ube3a^m-/p+^ mice adult mice (10-12 weeks of age), as single 150 ug ICV dose of RTR26183 was administered to mice by ICV injection. Brain regions (hippocampus, striatum and cortex) were harvested at 2 weeks post injection.

Ube3a^m-/p+^ and Ube3a^m+/p+^ animals were genotyped as previously described Jiang et al. 1998. [ICV injections were performed using a Hamilton microsyringe fitted with a 27 or 30-gauge needle, according to the method of Haley and McCormick. The needle was equipped with a polyethylene guard at 2.5 mm from the tip in order to limit its penetration into the brain. Mice were anesthetized using isoflurane anesthetic (1.5-4%). The needle tip was then inserted through the scalp and the skull into the right lateral ventricle, about 1 mm lateral and 1 mm caudal to bregma. Oligomer was given in a concentration of 20-40 mg/ml in a volume of 5 microliters in 0.9% saline. The needle was left in place for 10 seconds before removal. This procedure required no surgery or incision. Animals were warmed on heating pads until they recovered from the procedure. Brain tissue (right, frontal cortical region) was collected on dry ice or RNAlater for drug concentration analysis, Tau qPCR and protein measurement.

#### Non-human primate ASO dosing

Subjects were male cynomolgus monkeys weighing 2.5-3.4 kg at the start of the study. All animals got a polyurethane catheter implanted in the lumbar intrathecal space which was connected to a subcutaneous access port (Access Technologies) to allow for injection into the intrathecal space. Catheter placement at the lumbar site was confirmed by fluoroscopy. A contrasting agent (Omnipaque 240®, up to 2 mL) was injected into the catheter, and a dorsal image of the catheter was taken. Patency checks involving both injection of 0.2 mL of saline into lumbar ports and collection of CSF was performed weekly until completion of the study, except when dosing or CSF collections are scheduled. The reference or test item/article was administered to the appropriate animals using a syringe and needle via the subcutaneous access port connected to a catheter implanted in the intrathecal space at the lumbar level. Selected animals were dosed by a single dose on day 1 (all Groups) or two doses on days 1 and 15 (Groups 7 and 8 only). Animals were dosed in the prone position and maintained in the prone position for at least 30 min after the completion of dosing. Prior to dose administration, 1 mL of CSF was withdrawn; the first 0.2 mL of CSF was discarded. The dose volume for each animal was fixed at 1 mL and was given over 3 min. Following dose administration, the catheter was flushed with 0.5 mL/kg of artificial CSF (aCSF) given over a target of 1.5 min. Blood was collected by femoral venipuncture. Animals were fasted overnight before blood sampling. CSF samples were collected from the lumbar access port of all animals. The first 0.2 mL of CSF were discarded. After sample collection, the catheter was flushed with 0.2 mL of saline. Sample was centrifuged in a centrifuge set to maintain 4° C, for 5 min at 2000 x g within 1 hr of collection, and the supernatant was harvested and stored at - 80°C. All protocols and procedures involving the care and use of animals were reviewed and approved by the Institutional Animal Care and Use Committee of the testing facility where the study was conducted. The care and the use of animals were conducted according to the USA National Research Council and the Canadian Council on Animal Care (CCAC).

#### Tissue Readouts from in vivo experiments

##### 1) RNA analysis

Freshly frozen and weighed tissue samples (15–50 mg) were homogenized in 2 mL Precellys tube (Bertin Instruments) containing ceramic beads and 600 μL MagNaPure LC RNA Isolation Tissue buffer (Roche Life Science). The homogenate (400 μL) was then transferred to a MagNaPure 96 Processing Cartridge and RNA was purified using the MagNa Pure 96 with the kit Cellular RNA Large Volume Kit (05467535001) and using the protocol “RNA Tissue FF Standard LV 3.1” (Roche Life Science). Remaining lysate was stored for later analysis of ASO contend (see below).

RNA concentration of all samples was determined using the Eon Microplate Spectrophotometer (BioTek Instruments). Based on these concentrations, samples were diluted to 50 ng/μL and a number of random samples were checked for RNA quality using a Bianalyzer (Agilent). The purified RNA was further diluted between 1:10 to 1:100 (e.g. 5 μL RNA + 95 μL H2O) into a new RNA dilution plate (96-well PCR quality plate – Thermo Scientific #AB0900). The final in-well concentration of the diluted RNA was in the range of 0.2 to 5 ng/μL.

The RT-PCR reaction was carried out as a one-step RT-PCR reaction (XLT-Onestep, Quanta) on a ViiA 7 384-well Real-Time PCR System (Thermo-Fisher) running a standard 40 cycle program according to the manufacturer protocol. All reactions were carried out in duplicates and all samples from the same tissue were kept on the same QPCR plate. On each plate a standard curve was included with 2-fold serial dilution of RNA from the same tissue purified from saline treated control animals. Using the Quantstudio software (Applied Biosystems) RNA quantities were calculated based on the standard curve.

All reactions were carried out in duplicates and on each QPCR plate a standard curve was included with 2-fold serial dilution of purified RNA from treated control wells. Using the Quantstudio software (Applied Biosystems) RNA quantities were calculated based on the standard curve.

For analysis of cynomolgus tissue several housekeeping genes (GAPDH, POLR3F, UBC, and YWHAZ) were quantified from each sample and the three most stable genes were used for normalization in each tissue. The stability of HK genes was calculated using the method published by Vandesompele (Vandesompele et al., 2002). For assays, primers and probes used for housekeeping genes and target genes we used the following assays: *UBE3A* mRNA (Hs001666580_m1, Invitrogen), Gapdh (Mf04392546_g1, Invitrogen), Polr3F (Mf+2860939_m1, Invitrogen), Ywhaz (Mf029302410_m1, Invitrogen), Ubc (Mf02798368_m1, Invitrogen) and for *UBE3A-ATS* a custom assay from Integrated DNA technologies, Forward primer: 5’-CCA TCT CTG ATA AGG ATG ATT GAG G-3’, Reverse primer: 5’-GTT CAC AGG AGA CCA AAC AGA TA-3’ and a Zen™ dualquenced 5’-labeled PCR probe with the sequence: 5’-/56-FAM/TTT GGC TTG/zen/TTG ACA CCA GCA CAG /3IABkFQ/-3’

For analysis of RNA expression levels in mice we used following Taqman assays: *Ube3a* mRNA (Mm02580987_m1), *Ube3a-ATS* (Mm03455899_m1) and Gapdh (4351309) all from ThermoFisher. Quantities of *UBE3A* mRNA and *UBE3A-ATS* were normalized to quantities of Gapdh in the same sample.

##### 2) Allele specific digital PCR assay

*F*rom the available cohort of cynomolgus monkeys at Charles River Laboratories, Montreal, blood samples were drawn and DNA was extracted (Genewiz). Prior to paired end sequencing on an Illumina Hiseq platform, DNA from a 1.6 Mb region containing the *UBE3A* and *UBE3A-ATS* loci was enriched using Roche-Nimblegen gene enrichment library, all carried out at Genewiz. This identified a single nucleotide polymorphism (SNP) in an exonic region of UBE3A, Chr7: 3.339.686C→A. A total of 12 heterozygous UBE3A SNP carriers were used and assigned to the saline group (N=3) as well as d15 (N=3) and d29 (N=6).

A SNP digital PCR assay was set-up using Primetime probes, (Integrated DNA Technologies) covering the C→A SNP in position 3,359,686 of Chromosome 7. For detection of the C-allele we used a 5’FAM labeled probe (5’-CcgTCGTCtttTga-3’) (uppercase: locked nucleic acids, lower case: DNA) with a 3’ IowaBlack FQ quencher. For the A-allele we used 5’HEX labeled probe (5’-atCCgTCtTctttTga-3’) with 3’ Iowa Black FQ quencher. Amplification primers were place in exon3 and −4 of the cynomolgus monkey transcript AB179374, using forward primer (5’-GCTGGTTGTGGAGGAAATCT-3’) and reverse primer (5’GTAAATAGCCAGACCCAGTACTAT-3’). Droplets were generated on a BioRAD Automated Droplet generator, using the manufacturer’s protocol. Following droplet generation RT-PCR was run on a BioRad C1000 touch thermal cycler PCR machine, using the following program: 45°C 10 min, 40 cycles of 94°C for 30 s and 55°C for 30 s, 98°C for 10 min. Finally, droplets were analyzed on a Biorad QX200tm droplet reader and data analyzed using the Quantsoft Pro software. Linearity and specificity of the assay was confirmed by mixing different ratios of gBlocks/synthetic DNA (Integrated DNA technologies) covering the PCR amplicon and carrying either the C- or the A-allele.

To ascertain the parental origin of the two alleles in each animal we used the allele specific quantification from the midbrain. In midbrain, one allele was clearly dominating over the other allele also in treated animals. Midbrain is known to be a low exposure tissue for ASOs, also confirmed from our exposure data (Figure 4B panel A, moreover the qPCR quantification of *UBE3A-ATS* (Figure 4B panel B-C) confirmed that low exposure resulted in limited knock-down of *UBE3A-ATS*. This is highly indicative that in midbrain, the maternal imprinting is largely preserved due to low exposure and poor *UBE3A-ATS* knock-down (Figure 4B panel A), and as a consequence the maternal allele remains the most dominant allele after treatment with RO7248824. Moreover, in the absolute quantification of *UBE3A* mRNA expression there was no indication of exaggerated pharmacology in the midbrain. Hence, we could use the samples from the midbrain with least *UBE3A-ATS* knock-down to determine which allele in a given animal was of maternal (most expressed allele) and paternal (least expressed allele) origin.

##### 3) UBE3A protein analysis by Western blot

All tissues were placed in Precellys tubes after dissection and shock frozen. On the day of sample preparation, 200 μL of SDC lysis buffer were added to the tissues. The composition of the SDC lysis buffer for 30 mL was as follow: 300 mg of SDC (sodium deoxycholate), 85.99 mg of TCEP (Tris (2-carboxyethyl) phosphine hydrochloride), 112.21 mg of CAA (2-Chloroacetamide), 3 mL of Tris (pH = 8.5) and 3 tablets each of the protease/phosphatase inhibitors PhosStop and Complete Mini (Roche). Tissues were then lysed in Precellys Homogenizer Bertin (program 6 x 20s pulse, 6000 rpm) and placed on ice for 30 min. The supernatant was taken out and placed into 0.5 mL Eppendorf Protein Lobind tubes and boiled at 95°C for 5 min and then placed on ice. The supernatant was taken out and sonicated in Bioruptor (15×15s) and centrifuged. 5 μL of the supernatant was used for protein measurement by BCA Pierce and the rest was divided into aliquots and stored at −80°C. In preparation for the Western blot analysis 10 μL Loading buffer 4X and 4 μL NuPage Sample Reducing Agent 10X were added to 26 μL of each tissue sample. Water was then added to adjust to a final concentration of around 2.5 μg/μL of protein.

10 μL of tissue samples (25 μg of protein per well) were placed into each well of the Western blot gel (Biorad Criterion TGX Stain free 4-15% 18 wells; Cat. #5678085) and run in running buffer Tris-Glycine SDS (Invitrogen, Cat. #LC2675) under the following conditions: 60 V for 20 min and then 100 V for 100 min. The Western blot gels were analyzed with the software “Image Lab” on the Biorad imager and proteins transferred onto a nitrocellulose membrane with the device iBlot2 NC Regular Stacks (Invitrogen, Cat. #IB23001; program P0: step 1 - 20V/ 1 min, step 2 - 23 V/ 4 min, step 3 - 25 V/ 2 min). The membranes were blocked during 30 min in TBS/ 0.1% Tween20/ 3% milk at RT followed by incubation and shaking overnight at 4°C in 10 ml TBS/ 0.1% Tween20/ 3% milk containing 10 μl rabbit anti E6AP_Bethyl A300-352A. After washing three times for 5 to 10 min with TBS/ 0.1% Tween20/ 3% milk at RT the membranes were incubated and shaken for 90 min at RT in 10 ml TBS / 0.1% Tween20 / 3% milk including 2 μL goat anti-rabbit IgG (Jackson Laboratories, Cat #115-035-045) and washed 4 times for 5-10 min each with TBS/ 0.1% Tween20/ 3% milk at RT. For detection the membranes were shaken in 10 mL Lumi-light Western Blotting substrate (Roche, Cat. #12015200001) spiked in 1 mL SuperSignalTM West Femto Maximum Sensitivity Substrate (ThermoFischer, Cat. #34096) during 1 min and then analyzed on the Biorad imager.

Images of gels and blots were analyzed with the software “Image Lab”, which allowed to get a numerical value of the amount of protein in the gel or on the membrane. First, squares were drawn around one band per sample of the gel to select the total amount of protein on the gel. A second square was drawn next to the samples for subtraction of the background. The total protein values were called “gel”. The same was done for the nitrocellulose membrane and the UBE3A protein values were called “blot”. By calculating the “blot/gel” ratio (value on the blot per sample divided by the value on the gel) we got the proportion of UBE3A/total protein in the samples.

##### 4) Selective Monitoring Assay for Ube3a in mice

100 ug Tissue lysates prepared in SDC buffer for western blotting was used for digestion for SRM based LC-MS analysis. Samples were digested according to (*45*) with slight modifications. Briefly, SDC buffer containing samples were diluted in 4X in ddH2O and digested with 10 ug trypsin overnight at RT. Digested peptides were cleaned up in a 96 well Sep-Pak C18 96-well Plate: Waters. Cleaned peptides were dried in a speed vac (Thermo) for 4 hr.

1 ug of digested peptide was analyzed using LC-MS as described in (*8*). Skyline v 4.2 was used to generate a target peptide list based on shotgun MS analysis of Ube3a protein using a shared peptide between mice, cynomolgus monkey and human. Isotope-labeled peptide, containing either L-[U-13C, U-15N]R or L-[U-13C, U-15N]K, corresponding to the unique target peptide (VDPLETELGVK) was synthesized (Thermo) and their sequences confirmed by LC-MS/MS.

For MS analysis, digests were diluted to 200 ng/μL with 0.1% v/v formic acid, 2% v/v ACN containing the pooled isotope-labeled peptides at a final concentration of approximately 3 fmole/μL. SRM analyses were performed on an Ultimate RSLCnano LC coupled to a TSQ Quantiva triple quadrupole MS (Thermo Scientific). Samples (5 μl) were loaded at 3 μl/min for 6 min onto a 2 cm × 75 μm C18 trap column (Acclaim Pepmap 100, 3 μm, 300 Å, Thermo Scientific) in loading buffer (0.5% v/v formic acid, 2% v/v ACN). Peptides were then resolved on a 50 cm × 75 μm C18 analytical column with integrated electrospray emitter heated to 40 °C (Easy-SPRAY, 2 μm, 100 Å, Thermo Scientific) using the following gradient at a flow rate of 250 nL/min: 6 min, 98% buffer A (2% ACN, 0.1% formic acid), 2% buffer B (ACN + 0.1% formic acid); 90 min, 30% buffer B; 96 min, 60% buffer B; 98 min, 80% buffer B; 114 min, 80% buffer B; 115 min, 2% buffer B; 138 min, 2% buffer B. The TSQ Vantage was operated in 2 min retention time windows; cycle time, 1.5 s; spray voltage, 2600 V; collision gas pressure, 1.5 mTorr; Q1 and Q3 resolution, 0.7 FWHM; capillary temperature 240 °C.

Skyline version 4.2 was used for automated peak integration. In a small number of cases, the peak selection was manually corrected based on the isotope-labeled peptide elution time and transition ratio as well as the expected transition ratios based on the chromatogram library. A transition was removed from a peptide if obvious interference was identified (through comparison to the chromatogram library and the isotope-labeled peptides). Post data acquisition, total precursor peak areas were exported and further analysis was performed on Spotfire (Tibco). The endogenous peptide peak areas were corrected for using spiked-in heavy labelled peptides by applying a correction factor (Median heavy peptide intensity / heavy peptide area per sample).

### Mouse PKPD modeling

The relationship between *Ube3a-ATS* knock-down and the concentration was investigated, as well as the relationship between the three pharmacodynamic (PD) markers. All markers were measured on the various brain regions (hippocampus, striatum and cortex) and expressed as percentage of expression in comparison to vehicle-treated animals. The relationship between the brain concentrations and the *Ube3a-ATS* knock-down was assumed to be direct and without delay, as well as the relationship between the three PD markers. To note, the relationship between the *Ube3a* mRNA and the Ube3a protein was assumed to be compound independent. Therefore, data obtained from RTR26183 and RTR26235 were analyzed together in order to have enough information to derive a relationship between the two markers. All the analyses were performed in GraphPad Prism^®^ 8.4.2 using the nonlinear fit after log transformation of the data.

### Cynomolgus monkey PD modeling

Three PD markers were investigated. All markers were measured on various brain regions and expressed as percentage of expression in comparison to vehicle treated animals. To note, in cynomolgus monkeys the maternal allele is expressed, in contrast to AS patients. Therefore, in cynomolgus monkeys *UBE3A* mRNA and protein are expressed and at maximal effect expression is doubled.

In the PKPD study, the *UBE3A-ATS* knock-down is maximal for the entire study duration, in most brain regions except in the brainstem (midbrain, medulla and pons). The *UBE3A-ATS* knockdown is assumed to be the same across brain regions. The first time point in the study was at seven days and potential delay in *UBE3A-ATS* knock-down could not be observed. *UBE3A-ATS* knockdown was therefore fitted against the measured concentrations with a direct response model. The analysis of the *UBE3A* mRNA data is conducted on PCR data from the midbrain and cortex regions. The *UBE3A* mRNA elevation is maximal from the first to the last observation; delay could not be observed (first observation at day seven). Therefore, a direct response model is used to fit the *UBE3A* mRNA elevation against the *UBE3A-ATS* knock-down values.

At the protein level, a delay is observed before reaching the maximal elevation level. To fit the protein elevation an indirect response model is used and the protein half-life is estimated. At the protein level, only the observations in the cortex regions are used. We exclude the observations in the midbrain for two reasons 1) the PK variability is significant (data not shown) 2) protein and LNA content were measured from separate samples. Without the estimation of the PK interindividual variability the estimation of the protein half-life would be at least difficult and biased. An indirect response model is used and the protein half-life is estimated. The translation level of the UBE3A protein is assumed directly proportional to the *UBE3A* mRNA expression level. Therefore, if the *UBE3A* mRNA expression level is doubled, so is the protein.

Parameter estimation is performed using Monolix 2018R1 (The Monolix software, Analysis of mixed effects models, LIXOFT and INRIA, http://www.lixoft.com/) software using the linear approximation for the estimation of the Fisher Information Matrix.

## Acknowledgments

We would like to thank Children’s Hospital in Boston, under the leadership of Prof. Christopher Walsh, for recruitment of patients into the study; Harvard iPSC core facility and the team of Laurence Daheron for the reprogramming and quality control of hiPSC lines; Christoph Patsch for leading the generation of NSCs and supporting the screen and the Roche postdoctoral fellowship program for funding N.J.P., C.W., V.C., and M.T., and the Roche RiSE internship program for funding M.T. We thank Prof. Stormy Chamberlain for critically reviewing the manuscript and Silke Zimmermann, Laura Badi, Tony Kam-Thong, Roland Schmucki, Fabian Koechl, Nadia Anastasi, Hippolyte Gander and Nicole Hauser for the excellent technical support. We thank Anirvan Ghosh, Omar Khwaja, Kelly Bales, and Martin Ebeling for their support and critical input on the project.

